# Killing the enemy or making it your ally? The effects of thiostrepton on tumor associated macrophages

**DOI:** 10.1101/2023.03.23.533899

**Authors:** Diego A. Pereira-Martins, Jacobien R. Hilberink, Isabel Weinhäuser, Eduardo M. Rego, Gerwin Huls, Jan Jacob Schuringa

## Abstract

Attempts to convert a tumor supportive M2-polarized macrophage microenvironment into a tumor suppressive M1-polarized one have recently received considerable attention. However, the specificity of novel compounds, their mode-of-action, and efficacy on presumed target cells *in vivo* needs to be interpreted with caution.

## Results and discussion

It is increasingly becoming clear that the tumor immune microenvironment (TME) plays a critical role in tumor progression and drug resistance. For instance, we and others have identified that the presence of tumor supportive M2-polarized macrophages associates with the poorest prognosis in patients with acute myeloid leukemia (AML)^1-3^, in contrast to patients with a predominant M1-polarized TME which display good prognosis.^1^ Functionally, we could should show for the first time that while M1-polarized macrophages indeed suppressed leukemia development, M2-polarized macrophages were able to increase homing, self-renewal potential, alter the metabolome, and increase transformation capacity of leukemic blasts *in vivo*.^1^ Tools to convert a tumor supportive M2-polarized TME into a tumor suppressive one, preferably *in vivo*, are therefore of significant interest.

In the study published by Hu et al.,^4^ the authors performed a high-throughput phenotypic screen to identify compounds that repolarize tumor supportive M2 macrophages into anti-tumorigenic M1 macrophages and vice versa. The phenotypic changes between different macrophage subtypes were determined by a morphology-defined Z-score, as well as by transcriptome and flow cytometry analysis to confirm the respective macrophage subtype at baseline and after treatment. Based on these parameters, 37 out of 4000 FDA-approved drugs exhibited M2 to M1 repolarization capacity, including thiostrepton. While the data generated by this study significantly contributed to the scientific community, there are a few methodological and conceptual key points that we feel need to be addressed.

Firstly, we would like to raise some concerns about the method used to assess macrophage polarization, which sets the foundation of the study. In line with the authors, we observed a clear morphological distinction between M1 and M2 macrophages (Paper figure 1A – Comment figure 1A). Yet, flow cytometry-defined differences between macrophage subtypes were much more pronounced in our studies (Paper supplementary figure 1B – Comment figure 1B). Given the well-established literature on cytokine-mediated macrophage polarization^5,6^, we are not doubting that the desired macrophage subtype was generated at the initiation of the experiment. However, considering that the authors detected only minor changes by flow cytometry, we wonder whether the method applied to evaluate repolarization in the form it was executed was satisfactory. Since macrophages are known to be highly autofluorescent cells^7^ and knowing that autofluorescence can generate a false-positive signal^8^, insufficient blocking during the staining process could be the cause of such minor differences (Paper supplementary figure 1b). For this reason, we incubated detached macrophages 30 min prior to staining in PBS supplemented with 2 mM EDTA, 2% BSA, 0.02% NaN3 and 10% Fc block (more details are provided in the supplemental information file). While the authors detected a 1.2-fold increase in MFI for CD80 and CD86 in M1 compared to M2 macrophages, our results indicated a fold increase of 18 (M2a – IL4 polarized), -19.6 (M2d – IL6 polarized), and 3.87 (M2a), -28.7 (M2d), respectively. For CD163 and CD206 the authors measured a 1.5-fold increase in M2 compared to M1 macrophages, whereas we detected a fold increase of 2.3 (M2a) to 3 (M2d) and 5.5 (M2a) to 1.5 (M2d), respectively (Paper supplementary figure 1b – Comment figure 1B). Furthermore, we also observed a significant difference in percentage for M1 and M2 surface markers between different macrophage subtypes (Comment figure 1B).

**Figure 1.**
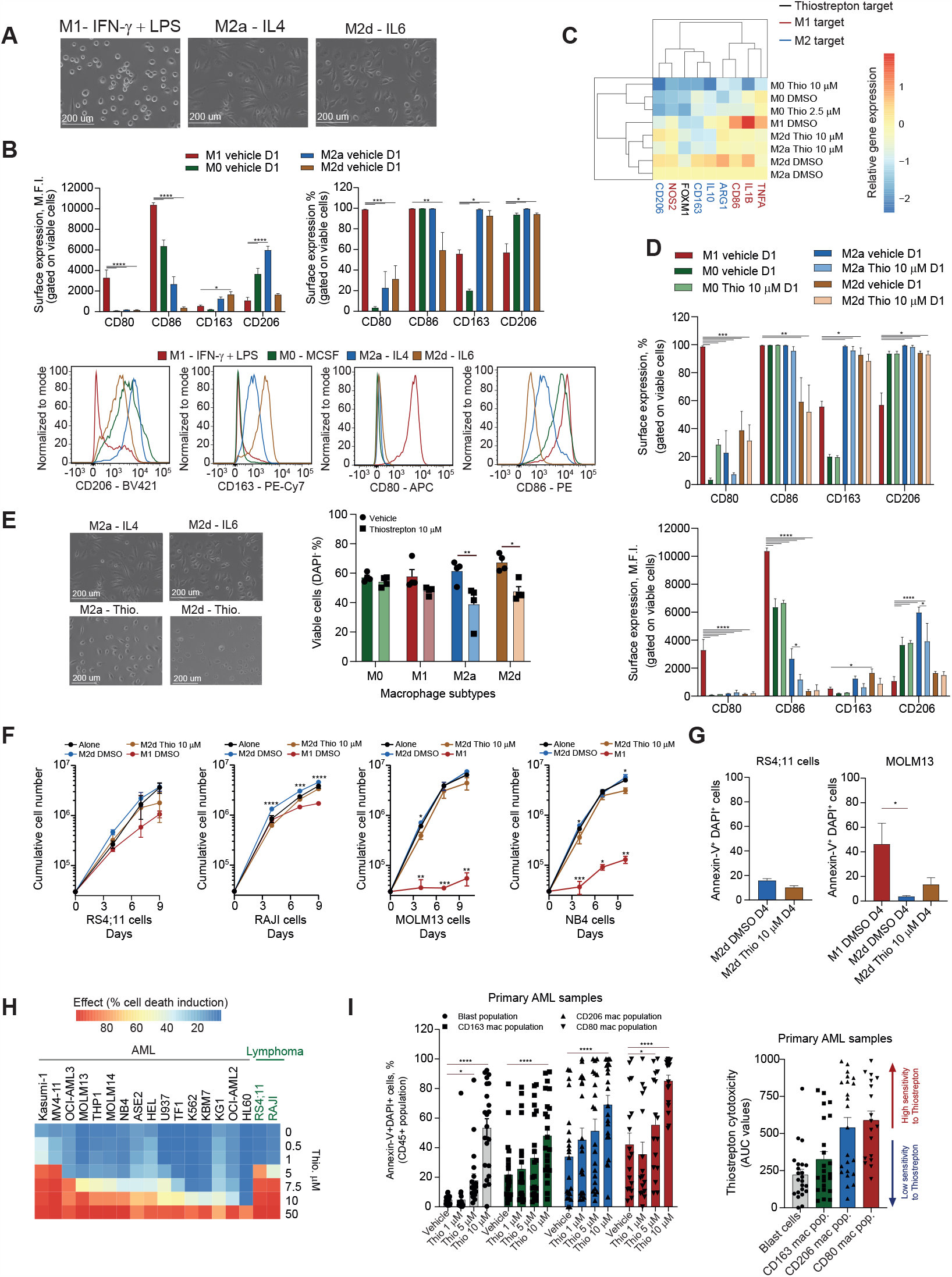
Effects of thiostrepton on macrophage polarization. Human macrophages were cultured for 24 h in the presence of LPS, IFNγ (for M1-polarization), IL-4 (M2a-polarization), and IL-6 (M2d-polarization). Representative pictures of M1, M2a and M2d macrophage morphology. Scale bar: 200 µm. (**B**) Bar plot comparing the levels (percentage, % and median fluorescence intensity, MFI) of M2- (CD163 and CD206) and M1- (CD80 and CD86) markers measured by FACS in healthy macrophages activated with respective cytokines for 24 h. M0-macrophages were generated with 50 ng/mL of MCSF (upper panels). Representative histograms of macrophage marker expression in different macrophage subtypes (lower panels). (**C**) Heatmap displaying the transcript levels of the indicated genes in M0-macrophages treated with 2.5 and 10 µM of Thiostrepton (DMSO was used as vehicle), and M2a- and M2d-macrophages treated with vehicle (DMSO) or Thiostrepton (10 µM) for 24 h. M1-macrophages were included as reference for the experiment. **p<0.01, ***p<0.001 indicate significance compared to all other groups. (**D**) Bar plot comparing the levels (MFI, upper panel; %, lower panel) of M2- (CD163 and CD206) and M1- (CD80 and CD86) markers measured by FACS in healthy activated M0, M1, M2a- and M2d-macrophages treated with vehicle and Thiostrepton (10 µM) for 24 h. (**E**) Cell morphologies of M2a and M2d activated macrophages treated with vehicle or Thiostrepton (10 µM) for 24 h. Scale bar: 200 µm. Bar plot comparing the viability (DAPI^-^ cells) of M0, M1, M2a, and M2d-macrophages treated with Thiostrepton and vehicle for 24 h. (**F**) Cumulative cell count of RS4;11, RAJI, MOLM13, and NB4 leukemic cell lines cultured on M1 and M2d macrophages (treated or not with Thiostrepton, 10 µM) for 9 days. (**G**) Percentage of apoptotic RS4;11 and MOLM13 cells after 4 days of culture on M1 and M2d treated with Thiostrepton (10 µM) or vehicle. (**H**) Effective dose 50 (ED^50^) of Thiostrepton in a panel of AML and B-ALL/lymphoma cell lines. (**I**) *Ex vivo* drug induced apoptosis of Thiostrepton in a set of 22 primary AML samples (Left panel). Area under the curve (AUC) values for Thiostrepton-induced cell death on blasts cells, M1, and M2 macrophages in AML samples. Cells were treated for 72 h, and apoptosis was evaluated by Annexin-V/DAPI staining by FACS. One-way (G left panel) or two-way (B-F and I) analysis of variance (ANOVA). *p<0.05, **p<0.01, ***p<0.001. (G right panel) Wilcoxon signed rank test (2-sided) *p<0.05. Source data are provided as a Source Data file.

Next, the authors evaluated the repolarization capacity of six M1 activating compounds that emerged from the compound screen by transcriptional and flow cytometry changes, including thiostrepton (Paper supplementary figure 3a and 3c; figure 4a and supplementary figure 5). Like the authors we focused our study on the M2 to M1 repolarization capacity of thiostrepton. When we evaluated the gene expression of the M1/M2 selected genes by the authors (primer sequences provided in the paper supplemental table 3) in M0 thiostrepton-treated macrophages, we could observe a decrease for some M2 markers such as *CD163* (p=0.03) and *IL10* (p=0.01) and the M1 markers *CD86* (p=0.04) and *TNFA* (p=0.04) when using the highest dose of thiostrepton (10 µM), but not for the dose that was mainly used by the authors (2.5 µM) (Comment figure 1C). Nevertheless, we could not detect a significant increase in M1-associated genes in any of the evaluated macrophage subtypes, although we did notice a reduction in the expression of thiostrepton target gene *FOXM1* (p=0.01) in both dosages, confirming the successful uptake of thiostrepton by the macrophages (Comment figure 1C). Additionally, the authors detected an MFI increase of approximately 20% for CD86 and a decrease of 10-20% for CD163 and CD206 respectively, while CD80 remained unchanged upon thiostrepton treatment (Paper supplementary figure 3c). Using M0-macrophages, we could not observe any significant change in percentage for the M1 markers CD80 and CD86, while the MFI of CD86 was decreased when using a dosage of 2.5 µM. Additionally, we also noted a decrease in MFI for the M2 marker CD206, which was dose dependent. (Comment supplemental figure 1A). Since for both gene expression and flow cytometry analysis, the treatment with the dose of 10 µM was the most effective, we expanded the analysis for other macrophage subtypes using the highest dose. In contrast with what was reported, even with the highest dose of thiostrepton we could not detect any significant change in MFI for CD80 and CD163, while M2a thiostrepton-treated macrophages presented a significant decrease in MFI of 55.7% and 34.4% for CD86 and CD206 respectively. No difference in percentage for any of the macrophage surface markers were observed in macrophage control versus thiostrepton treated macrophages (Comment figure 1D).

While we could not find compelling evidence for M2-M1 macrophage reprogramming by thiostrepton, we did note that the viability of M2 macrophages, but in fact also of most other cell types we investigated including leukemic cells themselves, appeared to be severely affected in the highest dosages (10 µM) (Comment figure 1E, H-I and comment supplemental figure 1A). This aligned with a more round-shaped morphology that was adapted by M2 macrophages upon thiostrepton treatment (Comment figure 1E), which might easily be misinterpreted as a repolarization towards M1-polarized macrophages (paper supplementary figure 1a).

In a next step the authors investigated functional changes in macrophage-supported tumor growth upon thiostrepton treatment. Short-term co-culture of B16F10 melanoma cells on B6 murine bone marrow-derived macrophages that were pretreated with thiostrepton led to a significant reduction in cell counts in a dose dependent manner and a significant increase of antibody-dependent cell-mediated phagocytosis (Paper figure 5e and supplementary figure 8b-c). Like the authors we co-cultured AML and lymphoma cell lines on human and murine thiostrepton-treated M2 macrophages for 12h (to observe the short-term mediated effects by thiostrepton-treated murine macrophages) and nine days (to observe the long-term effects of thiostrepton-treated human macrophages). Thiostrepton pretreatment of murine macrophages did not affect the leukemic growth in the short-term exposure (Comment supplementary figure 1B). In the human set, thiostrepton-treated macrophages barely impacted on leukemic cell growth, while the anti-proliferative effect of M1 macrophages was consistently observed (Comment figure 1F), albeit those lymphoid leukemic cells appeared to be less affected than myeloid leukemia cells. In line, no significant increase in apoptosis was observed when cells were grown on thiostrepton pretreated M2d macrophages, while apoptosis was significantly induced in AML lines on M1 macrophages (Comment figure 1G).

While we are aware that different cell lines have been used by the authors and that the anti-tumorigenic effects of M1 macrophages can vary between cell lines, as also observed in our data, our results indicate that M2 pretreated macrophages do not behave as M1-polarized macrophages. Our recommendation would be to always include M1-polarized macrophages in comparative studies. Although we are aware that the evaluation of macrophage repolarization goes beyond the detection of immunophenotypic/gene expression changes that are mainly associated with M1/M2 phenotypes, we support the idea that a macrophage repolarization would be associated with major phenotypic consequences, like an enhanced tumor-suppressive phenotype associated with M1-like macrophages. On a side note, one reason that could explain the difference in sensitivity to M1 anti-tumorigenic effects could be the metabolic differences across distinct cell lines. Previous studies as well as our unpublished observations showed that the secretion of lactate can repolarize M1 to M2 macrophages^9^ and our seahorse analysis indeed confirmed that lymphoma cell lines tended to be more glycolytic than AML cell lines (Comment supplementary figure 1C).

Next, we questioned whether thiostrepton is cytotoxic for cancer cells. When we evaluated the ED_50_ of AML and lymphoma cell lines treated with thiostrepton we determined an ED_50_ values, which ranged from 1.91 µM to 38.67 µM (n=16 AML cell lines; n=2 B-ALL/lymphoma cell lines, Comment figure 1H), indicating considerable sensitivity in tumor cells themselves as well. Similar observations were made by a recent publication showing thiostrepton mediated apoptosis in a panel of B-pre-ALL cell lines.^10^ Accordingly, thiostrepton also induced a significant increase in cell death in a dose dependent manner in *ex vivo* treated primary AML blast cells and in the monocytic subpopulation (with superior effects in this population), while healthy CD34^+^ cells remained largely unaffected (n=22 patients) (Comment figure 1I and supplementary figure 1D-E). In order to evaluate efficacy of thiostrepton treatment *in vivo* Hu et al.^4^ transplanted B6 and NSG mice with B16F10 and B lymphoma cells, respectively. Treatment of mice with thiostrepton showed a significant decrease in tumor burden (Paper figure 6a and supplementary figure 12a-b) and a slight increase of macrophage infiltration (Paper figure 6c-d and supplementary figure 12e). However, considering all the aforementioned points it cannot be excluded that the decrease in tumor burden is largely related to the cytotoxic effects of thiostrepton on the cancer cells themselves, or possibly on tumor-supportive M2-polarized macrophages.

Overall, the authors provide a great screening platform that allows the assessment of a large set of compounds on macrophage polarization. However, the morphological and flow-phenotypic definitions of macrophage subtypes can be experimentally challenging and should be performed with caution. Also, the cytotoxicity of identified macrophage-repolarizing compounds should be rigorously tested on leukemic cells themselves as well, as it is clearly relevant in the case of thiostrepton, in order to unravel the exact mode-of-action via which such compounds impact on leukemogenesis.

## Acknowledgements

This investigation was supported by Fundação de Amparo à Pesquisa do Estado de São Paulo (FAPESP, Grant #2013/08135-2). D.A.P-M. received a fellowship from FAPESP (Grant #2017/23117-1). I.W. received a fellowship from FAPESP (Grant #2015/09228-0). The authors would like to thank Dr. Emmanuel F. Griessinger for kindly providing the MS5, RS4;11, MOLM14 and RAJI cells used in the co-culture assays, as well as some of the leukemic lines used in the study.

I.W and D.A.P-M were sponsored by the Abel Tasman Talent Program (ATTP) of the Graduate School of Medical Sciences of the University of Groningen/University Medical Center Groningen (UG/UMCG), The Netherlands.

## Author contributions

D.A.P-M, J.R.H., I.W., and J.J.S. conceived and designed the study, performed experiments, analyzed, and interpreted data, performed the statistical analyses, and drafted the article. E.M.R. and G.H. provided patient samples and clinical data and reviewed the paper. All authors gave final approval of the submitted manuscript.

## Supplemental Materials and Methods

### EXPERIMENTAL MODEL AND SUBJECT DETAILS

#### Human sample collection and patient information

Mononuclear cells (MNCs) from AML patients were isolated via Ficoll (Sigma-Aldrich) separation and cryopreserved. Peripheral blood (PB) and bone marrow (BM) samples of AML patients were studied after informed consent and protocol approval by the Medical Ethical committee of the UMCG in accordance with the Declaration of Helsinki. Neonatal cord blood (CB) was obtained from healthy full-term pregnancies from the Obstetrics departments of the University Medical Center and Martini Hospital in Groningen, The Netherlands, after informed consent. The protocol was approved by the Medical Ethical Committee of the UMCG. Donors are informed about procedures and studies performed with CB by an information sheet that is read and signed by the donor, in line with regulations of the Medical Ethical Committee of the UMCG (protocol #NL43844.042.13). CB derived CD34^+^ cells were isolated by density gradient separation, followed by a hematopoietic progenitor magnetic associated cell sorting kit from Miltenyi Biotech (#130-046-702) according to the manufacturer’s instructions. All CD34^+^ healthy cells were pre-stimulated for 24-48 hrs prior to experimental use. CB derived cells were pre-stimulated with Stemline II hematopoietic medium (SigmaAldrich; #S0192), 1% penicillin/streptomycin (PS) supplemented with SCF (255-SC, Novus Biologicals), FLT3 ligand (Amgen) and N-plate (TPO) (Amgen) (all 100 ng/ml). Primary AMLs were grown on MS5 stromal cells with G-CSF (Amgen), N-Plate and IL-3, all 20 ng/ml.

#### Cell lines

All cell cultures were maintained in a humidified atmosphere at 37°C with 5% CO_2_. Mycoplasma contamination was routinely tested. All leukemia cell lines were authenticated by short tandem repeat analysis. MOLM14, RS4;11, RAJI and MS5 cells were kindly provided by Dr. Emmanuel F Griessinger (University Medical Centre Groningen, NL). MOLM14, RS4;11 and RAJI cells were culture in RPMI 1640 (Gibco, USA) and MS5 cells in IMDM (Gibco, USA) supplemented with 10% heat-inactivated fetal calf serum (HI-FCS) (Sigma-Aldrich, USA). ASE2 cells were kindly provided by Dr. M. Tomonaga (Dept. of Hematology, Nagasaki University School of Medicine, Nagasaki, Japan) and cultured in IMDM+20% HI-FCS + 5 units/mL of EPO (Amgen). KBM7 were kindly provided by the Brummelkamp lab (https://www.nki.nl/divisions/biochemistry/brummelkamp-t-group/) and cultured in RPMI + 20% HI-FCS. NB4 cells were kindly provided by Dr. Pier Paolo Pandolfi (Harvard Medical School, USA), and maintained in RPMI 1640 supplemented with 10% HI-FCS, L-glutamine (2 mM), sodium-pyruvate and penicillin/streptomycin (Invitrogen, USA). The THP1 (TIB-202™), K562 (CCL-243), MV4-11 (CRL-9591), and HL60 (CCL-240™) cell lines were obtained from the American Type Culture Collection and grown in RPMI 1640 (for THP1 and MV4-11) or IMDM (for K562 and HL60) with 10-20% HI-FCS (depending on the recommended dosage in the ATCC guidelines). All the other cell lines were obtained from the DSMZ-German Collection of Microorganisms and Cell Cultures. Mycoplasma contamination was routinely tested. All leukemia cell lines were authenticated by short tandem repeat analysis. Thiostrepton (CAS# 1393-48-2) was obtained from Calbiochem (Darmstadt, Germany).

#### Flow cytometry

Cryopreserved MNC fractions of AML patients were thawed, resuspended in newborn calf serum (NCS) supplemented with DNase I (20 Units/mL), 4 μM MgSO4 and heparin (5 Units/mL) and incubated at 37°C for 15 minutes (min). To analyze the myeloid fraction of the AML bulk sample after treatment with thiostrepton, 2.5×105 mononuclear cells were blocked with human FcR blocking reagent (Miltenyi Biotec) for 5 min and stained with the following antibodies: CD45-APC-Cy7, CD34-PE, CD14-PerCP, CD163-PE-Cy7, CD206-BV421, and CD80-APC for 20 min at 4°C. At the end, cells were washed and resuspended in IMDM medium supplemented with 20% of HI-FCS and Ca^2+^ buffer plus Annexin V FITC. A more mature myeloid population was detected based on the CD45 staining and inside this gate CD14 positive cells were then analyzed for their expression of the M1 marker CD80 and the M2 markers CD163 and CD206. Fluorescence was measured on the BD LSRII and analyzed using Flow Jo (Tree Star, Inc). For each sample a minimum of 5000 events were acquired inside the SSC-A^high^ CD45^high^ population.

#### Real Time quantitative polymerase chain reaction (RT-qPCR)

For quantitative RT-PCR, RNA (500 ng) was reverse transcribed using the iScript cDNA synthesis kit (Bio-Rad) and amplified using SsoAdvanced SYBR Green Supermix (Bio-Rad) on a CFX384 Touch Real-Time PCR Detection System (Bio-Rad). Primer sequences used for the analysis of M1 and M2-associated genes were obtained from the original paper of Hu et al 2021^1^ – Supplemental table 3 of the original paper.

#### Cord blood CD34 isolation

Peripheral blood mononuclear cells were isolated by a density gradient using Ficoll (Sigma-Aldrich) from cord blood. Mononuclear cells were washed once at 450g with PBS-EDTA (5 mM) and resuspended in 300 μL of PBS. Next, 100 μL of FcR blocking reagent and 100 μL of CD34 MicroBeads (Miltenyi Biotech) were added to the suspension and incubated for 30 min at 4°C. After incubation cells were washed for 10 min at 450g and resuspended in 2 mL of PBS–EDTA (5 mM). Cells were passed through a cell strainer (70 µm) and isolated by magnetic separation on the autoMACS (Program – Possedels, Miltenyi Biotech). The purity of the isolated cells was routinely evaluated by FACS and in the range of 85% to 95%.

#### Colony forming unit (CFU) assay

Cord blood isolated CD34^+^ cells were treated with thiostrepton (5 and 10 µM) and vehicle control (DMSO) and a total of 1×10^4^ cells were plated in semisolid methylcellulose medium supplemented with human cytokines MethoCult™ H4435 (StemCell). Colonies were detected after 14 days and scored.

#### Human macrophage generation

Mononuclear cells were isolated by a density gradient using Ficoll (Sigma-Aldrich) from healthy donors or allogeneic donors. Next, 2.5×10^6^ or 5×10^6^ mononuclear cells were seeded into 12 or 6-well plates and incubated for 3h at 37ºC in RPMI 1640 medium supplemented with 10% HI-FCS, 10% HI- and filtered human serum (AB serum, Thermofisher) and 1% penicillin and streptomycin (PS). After 2h of incubation, the non-adherent cell fraction was removed, and the new RPMI medium supplemented with 10% HI-FCS, 10% HI-AB serum and 1% penicillin and streptomycin was added. Additionally, 50 ng/mL of GM-CSF (Prepotech) or M-CSF (Prepotech/Immunotools) growth factors were added to the medium to generate pre-orientated M1 and M2 macrophages, respectively. Monocytes were differentiated into macrophages over a time span of 7 days and at day 3 half of the medium was renewed.

#### Murine macrophage generation

Murine macrophages were derived from NSG (NOD.Cg-Prkdcscid ll2rgtm1Wjl/SzJ) mice that were purchased from the Central Animal Facility breeding facility within the UMCG. All *in vivo* studies were conducted with ethical approval from the UMCG. Mice were anesthetized with an overdose of ketamine/xylazine solution and euthanatized by cervical dislocation to collect femur and tibia. Next the epiphyses were cut off and a syringe of 25 G filled with PBS (1% HI-FCS) was used to flush the BM onto a 70 µm cell strainer placed on a 50 ml tube. Red blood cells were lysed for 10 min at 4ºC and washed with PBS. Three million BM mononuclear cells were seeded in a 100×20 mm petri dish and cultured in RPMI 1640 medium supplemented with 10% HI-FCS, 15% of L929 supernatant and 1% PS for 7 days. At day 3 half of the medium was renewed. For co-culture experiments, murine macrophages were stained with CD45.1-PE and Gr1 APC to exclude human cells from the murine populations, and analyzed by flow cytometry.

#### Macrophage polarization

At day 7 human macrophages were washed with PBS and new RPMI 1640 medium supplemented with 10% HI-FCS was added. GM-CSF cultured macrophages were polarized to M1 macrophages with 20 ng/ml IFN-y (Prepotech) and 100 ng/ml LPS (Sigma-Aldrich), while M-CSF cultured macrophages were polarized to M2d macrophages with 10 ng/ml M-CSF and 20 ng/ml IL-6 or M2a with 10 ng/mL M-CSF and 20 ng/mL IL-4 (Prepotech). To generate M0 macrophages, M-CSF cultured macrophages were kept with 50 ng/mL of M-CSF in the medium.

#### Flow cytometry staining of macrophages

After polarization, macrophages are detached with TrypLE (ThermoFisher) according to manufacturer’s instructions. A total of 1×10^5^ cells were washed in PBS and resuspended in PBS (2 mM EDTA, 2% BSA, 0.02% NaN_3_) and 10 μL FcR blocking reagent. Macrophages were incubated 30 min at 4°C. Staining was performed after that with the M1 and M2 markers for 20 min at 4°C. Cells were washed and resuspended in PBS 1X plus a viability marker (Annexin-V or DAPI, depending on the panel combination of markers). Fluorescence was measured on the BD LSRII and analyzed using Flow Jo (Tree Star, Inc). For each sample a minimum of 10 000 viable cells (DAPI negative events) were acquired.

#### *In vitro* AML cell line proliferation on macrophages

Human macrophages were generated and polarized as described in the sections “Macrophage generation” and “Macrophage polarization”. A total of 3×10^4^/mL AML cells (NB4 and MOLM13) or lymphoma cells (RS4;11 and RAJI) were put in co-culture with M1 and M2d macrophages (treated or not with thiostrepton 10 µM). The co-cultures were performed in RPMI 1640 supplemented with 10% HI-FCS and 1% PS. The proliferation rate was assessed by daily cell counts with a hemocytometer and by FACS (LSRII) for a total period of 10 days.

#### Apoptosis assay

For the apoptosis analysis a minimum of 1×10^5^ AML cells were stained after 4 days of culture on M1, M2d macrophages (treated or not with Thiostrepton 10 µM). The apoptosis rate was determined using the Annexin V-FITC antibody and DAPI as a viability dye, in the presence of Ca^2+^ binding buffer. All specimens were acquired by flow cytometry (BD LSRII) and analyzed with the FlowJo software (Treestar, Inc., USA). All experiments were performed in triplicate and for each sample a minimum of 10 000 events were acquired.

#### Oxygen consumption (OCR) and extracellular acidification rate (ECAR) measurements

Oxygen consumption rate (OCR) and Extra Cellular Acidification Rate (ECAR) were measured using Seahorse XF96 analyzer (Seahorse Bioscience, Agilent, US) at 37 °C. For AML cell lines, 1×10^5^ viable cells (DAPI^-^) were seeded per well in poly-L-lysine (Sigma-Aldrich) coated Seahorse XF96 plates in 180 μL XF Assay Medium (Modified DMEM, Seahorse Bioscience), respectively. For OCR measurements, XF Assay Medium was supplemented with 10 mM Glucose and 2.5 µM oligomycin A (Port A), 2.5 µM FCCP (carbonyl cyanide-4-(trifluoromethoxy) phenylhydrazone) (Port B) and 2 µM antimycin A together with 2 µM Rotenone (Port C) were sequentially injected in 20 µL volume to measure basal and maximal OCR levels (all reagents from Sigma-Aldrich). For ECAR measurements, Glucose-free XF Assay medium was added to the cells and 10 mM Glucose (Port A), 2.5 µM oligomycin A (Port B) and 100 mM 2-deoxy-D-glucose (Port C) (all reagents from Sigma-Aldrich). All XF96 protocols consisted of 4 times mix (2 min) and measurement (2 min) cycles, allowing for determination of OCR at basal and also in between injections. Both basal and maximal OCR levels were calculated by assessing metabolic response of the cells in accordance with the manufacturer’s suggestions. The OCR measurements were normalized to the viable number of cells used for the assay.

#### Lactate secretion measurements

Lactate concentration was determined from the cell culture medium by monitoring NAD(P)H increase occurring during specific enzymatic reactions for each metabolite at 340 nm wavelength, as described elsewhere.^2^ Briefly, extracellular lactate concentrations were determined by the lactate dehydrogenase (LDH) enzymatic reaction in the cell culture medium taken at 0 hour and after co-culture of tumor cells with MS-5 or M2d macrophages, for 48 hours incubation. Cells were seeded in liquid culture after co-cultures, and culture medium was taken after 24 hours. Extracellular lactate was converted by L-Lactic Dehydrogenase (LDH, Sigma Aldrich) reaction in freshly prepared 25 mM NAD^+^ and 87.7 U/mL LDH in 0.4 M hydrazine (Sigma Aldrich)/0.5M glycine assay buffer (pH 9). 20 µl samples (diluted according to standard curve) and Sodium L-lactate (Sigma Aldrich) standards were pipetted into 130 µL reagent mix in 96 wells plate format and the reaction was carried out for 30 min at 37 °C.

### QUANTIFICATION AND STATISTICAL ANALYSIS

Statistical significance of the difference between multiple experimental groups was analyzed by one-way or two-way ANOVA (Kruskal Wallis test with posthoc Dunn analysis) using GraphPad PRISM 8 software or two-tailed paired or unpaired student’s t-test using excel and GraphPad PRISM 8 software. P-values are indicated in Figure. legends. For Fig. 1G a Wilcoxon signed rank test (2-sided) was used. P-values are indicated in Figure.

### Data availability

The source data underlying Figs 1B-I, and Supplementary Figs 1A-E are provided as a Source Data file All data are also available from the corresponding author on request.

## Resources table

| REAGENT or RESOURCE | SOURCE | IDENTIFIER |
| --- | --- | --- |
| **Antibodies** |  |  |
| Anti-Human CD45 FITC (1:100 dilution) | BioLegend | 368508<br>Clone: 2D1 |
| Anti-Human CD45 APC-Cy7 (1:100 dilution) | Biolegend | 368516<br>Clone: 2D1 |
| Anti-Human CD14 PerCP (1:100 dilution) | BioLegend | 325632<br>Clone: HCD14 |
| Anti-Human CD34 PE (1:100 dilution) | BD Biosciences | 348057<br>Clone: 8G12 |
| Anti-Human CD117 APC (1:100 dilution) | BD Biosciences | 550412<br>Clone: YB5.B8 |
| Anti-Human CD163 PE-Cy7 (1:100 dilution) | BioLegend | 333614<br>Clone: GHI/61 |
| Anti-Human CD206 BV421 (1:100 dilution) | BD Biosciences | 564062<br>Clone: 19.2 |
| Anti-Human CD80 APC (1:100 dilution) | BioLegend | 305220<br>Clone: 2D10 |
| Anti-Human CD86 PE (1:100 dilution) | BloLegend | 305406<br>Clone: IT2.2 |
| Annexin FITC (1:200) | Immunotools | 31490013X2 |
| Annexin APC (1:200) | Immunotools | 31490016X2 |
| Anti-mouse Gr1 APC (1:100 dilution) | BioLegend | 108412<br>Clone: RB6-8C5 |
| Anti-mouse CD45.1 PE (1:100 dilution) | BioLegend | 110708<br>Clone: A20 |
| **Bacterial and virus strains** |  |  |
| **Biological samples** |  |  |
| Human AML blast cells and healthy cord blood isolated CD34<sup>+</sup> cells | UMCG | Ethical committee<br>NL43844.042.13 |
| Murine bone marrow mononuclear cells | USP | Ethical committee<br>#176/2015 |
| Human peripheral blood mononuclear cells | UMCG | Ethical committee<br>NL43844.042.13 |
| **Chemicals, peptides, and recombinant proteins** |  |  |
| 4',6-diamidino-2-phenylindole | Sigma-Aldrich | 28718-90-3 |
| DNase I | Roche | 11284932001 |
| MgSO<sub>4</sub> | Sigma-Aldrich | M7506 |
| Heparin | Sigma-Aldrich | 60800-63-7 |
| Thiostrepton | Calbiochem | 1393-48-2 |
| Human Granulocyte-macrophage colony-stimulating factor | Peprtech | 300-03 |
| Human Macrophage Colony-Stimulating Factor | Immunotools | 11343117 |
| Human Interleukin 6 | Peprtech | 200-06 |
| Human Interleukin 4 | Peprtech | 200-04 |
| Human Interferon Gamma | Peprtech | 300-02 |
| Human Interleukin 3 | Peprtech | 200-03 |
| Human Granulocyte colony-stimulating factor | Peprtech | 300-23 |
| Human Thrombopoietin | Amgen |  |
| Human Erythropoietin | Amgen |  |
| Lipopolysaccharides | Sigma-Aldrich | L4391 |
| β-mercaptoethanol | Merck Sharp & Dohme BV | 60-24-2 |
| SsoAdvanced Universal SYBR® Green Supermix | BioRad | 1725274 |
| iScript cDNA synthesis Kit | BioRad | 1708891BUN |
| **Critical commercial assays** |  |  |
| Seahorse XFe96 Flux Analyzer | Agilent |  |
| **Deposited data** |  |  |
| **Experimental models: Cell lines** |  |  |
| MOLM13 | DSMZ | ACC 554 |
| MV4-11 | ATCC | CRL-9591™ |
| HL60 | ATCC | CCL-240™ |
| OCI-AML3 | DSMZ | ACC 582 |
| NB4 | Harvard Medical School | Prof. Pier Paolo Pandolfi |
| MS-5 | UMCG | Dr. Emmanuel F Griessinger |
| RAJI | UMCG | Dr. Emmanuel F Griessinger |
| RS4;11 | UMCG | Dr. Emmanuel F Griessinger |
| MOLM14 | UMCG | Dr. Emmanuel F Griessinger |
| ASE2 | Nagasaki University School of Medicine | Dr. M. Tomonaga |
| KBM7 | Brummelkamp lab | Dr. Thijn Brummelkamp |
| Kasumi-1 | DSMZ | ACC 220 |
| HEL | DSMZ | ACC 11 |
| KG1 | DSMZ | ACC 14 |
| TF1 | DSMZ | ACC 334 |
| U937 | DSMZ | ACC 5 |
| THP1 | ATCC | TIB-202™ |
| K562 | ATCC | CCL-243 |
| OCI-AML2 | DSMZ | ACC 99 |
| L929* | ATCC | CCL-1™ |
| **Experimental models: Organisms/strains** |  |  |
| NSG (NOD.Cg-Prkdcscid Il2rgtm1Wjl/SzJ) | Jackson Laboratory | 005557<br>RRID:<br>IMSR_JAX:005557 |
| **Oligonucleotides** |  |  |
| Primer sequences were available at supplemental table 3 of the main manuscript – Hu et al., 2021 |  |  |
| **Recombinant DNA** |  |  |
| **Software and algorithms** |  |  |
| FlowJo v10.0.6 | Treestar | http://www.flowjo.com/ |
| Prism 9 | GraphPad | http://www.graphpad.com/ |
| SPSS Statistical package 19.1 | IBM | https://www.ibm.com/ |
| Wave | Agilent | https://www.agilent.com/ |
| RStudio | CRAN | www.r-project.org |
| **Other** |  |  |
| FcR Blocking reagent, human | Miltenyi Biotec | 130-059-901<br>RRID: AB_2892112 |
| CD34 MicroBeads Kit UltraPure, Human | Miltenyi Biotec | 130-100-453 |
| TrypLE™ Express Enzyme | Thermo Fischer Scientific | 12604021 |
| MethoCult™ | Stemcell | H4435 |

**Supplemental figure 1.**
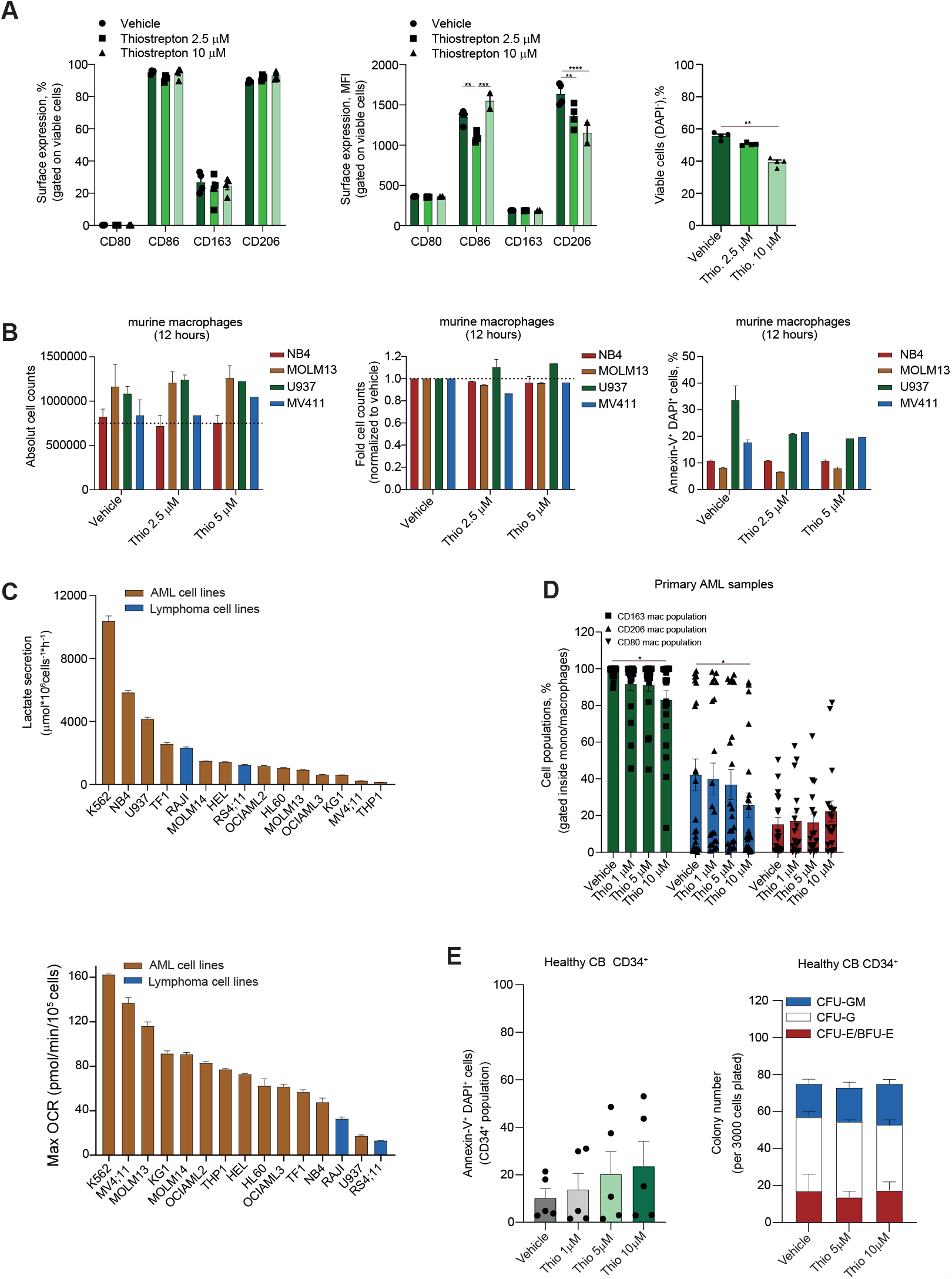
(**A**) Bar plot comparing the levels (left panel - percentage, % and middle panel - median fluorescence intensity, MFI) of M2- (CD163 and CD206) and M1- (CD80 and CD86) markers measured by flow cytometry in Peripheral blood isolated healthy macrophages cultured in the presence of 50 ng/mL recombinant human MCSF (M0-macrophages) for 7 days and treated in the absence of cytokines with thiostrepton for 24 h. Right panel shows the Bar plot comparing the viability (DAPI-cells) of M0-macrophages treated with Thiostrepton at 2.5 and 10 µM and vehicle for 24 h. Representative histograms of macrophage marker expression in different macrophage subtypes (lower panels). (**B**) Cumulative cell counts (left panel), fold of proliferation relative to vehicle (middle panel) and apoptosis levels (evaluated by Annexin-V/DAPI staining) of human leukemic cells co-cultured for 12 h on murine bone-marrow isolated macrophages. Macrophages were pre-treated with vehicle (DMSO) or Thiostrepton (2.5 and 5 µM) prior to the co-culture (24 hours). For flow-cytometry evaluations, cells were stained with human CD45-FITC to exclude cross-contamination with floating murine macrophages. (**C**) Lactate secretion per 1 × 106 cells per hour (upper panel) and maximum oxygen consumption rate (Max OCR, lower panel) in leukemic cells (n = 4 measured in quadruplicates). (**D**) Bar plot comparing the levels macrophages expressing CD163, CD206, and CD80 measured by flow cytometry in ex vivo AML samples treated with Thiostrepton (doses: 1, 5, and 10 µM) for 72 h. (**E**) Ex vivo drug induced apoptosis of Thiostrepton in a set of 05 independent healthy CD34^+^ cells isolated from cord blood samples (Left panel) and colony formation capacity (right panel). Cells were treated for 72 h, and apoptosis was evaluated by Annexin-V/DAPI staining by flow cytometry. Clonogenic assay was performed in methylcellulose medium supplemented with human cytokines. Colonies were scored after 14 days of plating. One-way or two-way analysis of variance (ANOVA) were used for statistical analysis. *p<0.05, **p<0.01, ***p<0.001. Source data are provided as a Source Data file.

